# Dronedarone hydrochloride reverses obesity-related metabolic syndrome while preserving skeletal muscle mass

**DOI:** 10.64898/2026.07.20.739085

**Authors:** Jiaxin Lei, Xiaoxun Zhang, Xinyu Cao, Zhixian Zhu, Fei Ye, Ziqian Xu, Wenrong Su, Xinyue Zeng, Zuzhi Xu, Jingdong Zhao, Siyi Jiang, Nan Zhao, Huaizheng Liu, Yan Lu, Chuanzheng Sun, Jin Chai

**Affiliations:** Department of Gastroenterology, The First Affiliated Hospital (Southwest Hospital) of Third Military Medical University (Army Medical University), Chongqing, China; Institute of Digestive Diseases of PLA, Third Military Medical University (Army Medical University), Chongqing, China; Cholestatic Liver Diseases Center, The First Affiliated Hospital (Southwest Hospital) of Third Military Medical University (Army Medical University), Chongqing, China; Metabolic Dysfunction-Associated Steatotic Liver Disease (MASLD) Medical Research Center, The First Affiliated Hospital (Southwest Hospital) of Third Military Medical University (Army Medical University), Chongqing, China; Department of Infectious Diseases, Xiangya Hospital, Central South University, Changsha, Hunan, China; Department of Emergency, The Third Xiangya Hospital, Central South University, Changsha, China; Institute of Metabolism and Regenerative Medicine, Shanghai Sixth People’s Hospital Affiliated to Shanghai Jiao Tong University School of Medicine, Shanghai, China

**Keywords:** dronedarone hydrochloride, obesity

## Abstract

Obesity-driven metabolic syndrome poses a critical global threat, yet standard therapies like GLP-1 receptor agonists trigger substantial lean mass wasting, with muscle loss accounting for up to 40% of reduced weight. Here we identify a non-canonical metabolic application for dronedarone hydrochloride, an anti-arrhythmic benzofuran derivative. In diet-induced and ob/ob obese mice, short-term dronedarone hydrochloride administration dose-dependently reduces food intake, clears visceral and subcutaneous adiposity, and reverses steatohepatitis. Head-to-head trials show that dronedarone hydrochloride achieves glycemic control and fat clearance non-inferior to semaglutide, tirzepatide, and empagliflozin, but uniquely and completely preserves skeletal muscle mass. Mechanistically, dronedarone hydrochloride operates independently of central hypothalamic appetite-regulating neuropeptides and the peripheral leptin pathway. By decoupling fat reduction from sarcopenia, our findings establish dronedarone hydrochloride as a muscle-sparing therapeutic candidate for metabolic syndrome.

## INTRODUCTION

Obesity has escalated into a defining global pandemic of the 21st century, with the age-standardized prevalence in adults more than doubling since 1990 and surpassing one billion people worldwide [1, 2]. Characterized by the excessive accumulation of visceral adipose tissue, obesity drives systemic chronic inflammation and mitochondrial dysfunction [2–4]. This pathological cascade serves as the core mechanism triggering insulin resistance and driving the onset of metabolic syndrome, a cluster of disorders including type 2 diabetes, metabolic dysfunction-associated steatohepatitis, hypertension, and dyslipidemia [5]. Collectively, these multi-organ metabolic abnormalities act synergistically to accelerate atherosclerosis, significantly elevating cardiovascular mortality, inducing musculoskeletal damage, and raising the risk of various malignancies [6].

However, existing therapeutic modalities remain tightly constrained by critical clinical limitations and paradoxes. Although new-generation GLP-1 receptor agonists (e.g., semaglutide, tirzepatide) achieve profound weight loss and improve cardiovascular outcomes, up to 50% of patients experience gastrointestinal adverse events, and 4.5-7.1% discontinue treatment due to intolerance [7–9]. Crucially, approximately 25-40% of the weight lost under these therapies is lean mass rather than fat mass, precipitating a dangerous risk of sarcopenia and a subsequent decline in basal metabolic rate that contributes to rapid weight regain upon discontinuation [10]. SGLT-2 inhibitors such as empagliflozin offer excellent glycemic control but exhibit relatively limited efficacy in driving substantial, direct lipid-lowering and visceral fat clearance. Furthermore, invasive options like bariatric surgery are irreversible, carry an inherent procedural mortality risk, and produce severe long-term bone mineral density loss and skeletal muscle wasting. Thus, there is an urgent, unfulfilled clinical need to develop novel metabolic intervention strategies that can aggressively clear pathological fat while uniquely preserving skeletal muscle mass [11].

Dronedarone hydrochloride, a non-iodinated benzofuran derivative approved by the United States Food and Drug Administration in 2009 for maintaining sinus rhythm in atrial fibrillation, presents an intriguing candidate for drug repositioning [12]. Structurally analogous to amiodarone, dronedarone hydrochloride inhibits multiple ion currents and β1-adrenergic receptors, yet the strategic deletion of the iodine moiety eliminates the severe thyroid, pulmonary, and hepatic toxicities that limit long-term amiodarone use [12]. While landmark clinical trials-including DAFNE, ATHENA, and DIONYSOS-have robustly established its safety and efficacy profiles in cardiovascular medicine [13–15], the potential application of dronedarone hydrochloride in managing systemic energy homeostasis and metabolic disorders remains entirely unexplored.

This study uncovered an unconventional mechanism underlying dronedarone hydrochloride’s anti-obesity and metabolic protective effects. Short-term treatment with the compound reversed diet-induced insulin resistance and multi-organ metabolic dysfunction. In direct comparisons, dronedarone hydrochloride alleviated adipocyte hypertrophy and hepatic lipid accumulation with efficacy comparable to first-line therapies (semaglutide, tirzepatide, empagliflozin). Unlike these agents, it did not reduce lean mass. By separating fat mass reduction from lean tissue loss, our data support dronedarone hydrochloride as a muscle-sparing candidate for long-term intervention in obesity-related metabolic syndrome.

## MATERIALS AND METHODS

### Animal experiments and procedures

All mouse colonies were maintained at the Animal Core Facility of the First Affiliated Hospital of Army Medical University. The animal studies were approved by the Army Medical University Institutional Animal Care and Use Committee (Approval No. AMUWEC20232171). We purchased a batch of 6-week-old C57BL/6J and ob/ob (Lep KO) male mice from Caygen Biotechnology Company. For the C57BL/6J mice, we randomly assigned them into several groups and fed them a high-fat diet(CAT#D12492, Research Diets) or NCD (CAT#D12450J, Research Diets) for 12 weeks. All of the mice were maintained under a 12 h light/dark cycle at a constant temperature of 22 °C and relative humidity of ∼50%, with food and water provided in a SPF level animal room. After reaching the designated feeding time, administered different treatments to the mice and recorded the daily body weight and food intake. As for the ob/ob mice, we administered either the drug or a control solvent directly for 3 weeks. The solvent used for dissolving dronedarone hydrochloride is 5% Dimethyl Sulfoxide(Solarbio, Cat#D8371, China) and 6% Polyethylene glycol 6000(Aladdin, P103725-500g, China). Semaglutide (TargetMol, Cat#910463-68-2, China), Tirzepatide (TargetMol, Cat#2023788-19-2, China) and Empagliflozin (TargetMol, Cat#864070-44-0, China) are prepared according to the instructions. Once the intervention was completed, the mice were euthanized. Liver, adipose tissue, gastrocnemius and serum were collected. The levels of serum alanine transaminase (ALT) and total cholesterol (TC) were measured at the Clinical Laboratory Department of the Second Affiliated Hospital of Army Medical University.

### Oral Glucose Tolerance Test (OGTT)

The mice were fasted overnight (12 hours) prior to the test, whilst ensuring they had access to water. Weigh all the mice and mark their tails with a permanent marker so that they can be easily distinguished. Then, mice were gavaged with 20% glucose solution (1.5 g/kg). Blood samples were collected from the tail vein at thirty-minute intervals (0, 30, 60, 90 and 120 min), with blood glucose levels recorded for each animal.

### Insulin Tolerance Test (ITT)

Before insulin injection, fast the mice for 4 hours, whilst ensuring they have access to water. Weigh the mice, mark their tails and take blood samples from the tail vein to measure baseline blood glucose levels. Then, mice were intraperitoneally injected with insulin (0.75 U/kg) (Actrapid, Novo Nordisk, Bagsværd, Denmark). By detecting the tail vein, we recorded the blood glucose levels of mice at 0, 15, 30, 45 and 60 minutes.

### Hematoxylin and eosin (H&E)

Mouse liver, eWAT, and iWAT tissue samples were immersed in 4% PFA for 48 hour before paraffin-embedded. Then we sliced the samples into 5-µm-thick sections. The sample sections were stained with H&E (Solarbio; Cat#G1121; China) following the manufacturer’s instructions. Expert pathologists were invited to evaluate these sections using the NAS scores in a blinded manner.

### Micro-CT analysis of mouse body composition

Micro-computed tomography (micro-CT) is a non-invasive high-resolution imaging technique. This device captures multi-angle X-ray two-dimensional projection images of samples and reconstructs a three-dimensional ray density dataset. By leveraging density differences among various tissues, algorithms can perform quantitative in situ segmentation and calculate the volume of adipose tissue. The mice were administered gas anesthesia to facilitate in vivo imaging. The fixed region was scanned (upper boundary: superior edge of the L1 vertebra; lower boundary: highest point of the ilium), with a uniform number of slice layers maintained. Finally, we analyzed the scan results to obtain quantitative adipose tissue data.

### Quantitative real-time PCR

Total RNA was isolated from snap-frozen mouse hypothalamus tissue using the RNAiso Plus(Takara Bio, Kyoto, Japan). We used HisyGo RT Red SuperMix (Vazyme Bio, Nanjing, China, Cat#RT101-01) to reverse transcribe total RNA (1 µg) to cDNA. The expression of mRNA was assessed using SYBR Green r eal-time PCR(Vazyme Bio, Nanjing, China, Cat#Q412-02). Gene-specific primer sequences were these (Agrp-F: 5′- CTTTGGCGGAGGTGCTAGAT-3′, R: 5′-A GGACTCGTGCAGCCTTACAC-3′; Npy-F: 5′-CTCCGCTCTGCGACACTACA-3′, R: 5′-AATCAGTGTCTCAGGGCTGGA-3′; Pomc-F: 5′-CAGTGCCAGGACCTC ACCA-3′, R: 5′-AGCGAGAGGTCGAGTTTGCA-3′; Cart-F: 5′-AAGTCCCCAT GTGTGACGCT-3′, R: 5′-GACAGTCACACAGCTTCCCGA-3′). Transcript levels were normalized to 18S gene and relative gene expression was calculated using 2^-ΔΔCt^ method.

### Statistical analysis

Data are presented as the mean ± standard deviation. Statistical analyses were performed using the SPSS statistics software. The comparisons between the two groups were analyzed using an unpaired two-tailed Student’s t-test, Welch’s t test, or one-tailed Mann-Whitney U test. One-way analysis of variance (ANOVA) was applied to determine the comparisons involving multiple groups. Statistical significance was set at p < 0.05.

## RESULTS

### Intraperitoneal injection of dronedarone hydrochloride can treat obesity-related metabolic disorders in mice with a high-fat diet

To investigate the role of dronedarone hydrochloride in obesity and metabolic disorders, we established an obese mouse model which was induced by a high-fat diet **(Figure 1A)**. C57BL/6J mice were randomly divided into four groups and fed either NCD or HFD diets for 12 weeks. Subsequently, we administered low-dose dronedarone hydrochloride (10mg/kg), or high-dose dronedarone hydrochloride (30mg/kg) intraperitoneally to obese mice every day for two weeks, while monitoring their body weight and daily food intake, and plotted the data as a curve chart **(Figure 1B-C)**. We found that dronedarone hydrochloride significantly influenced body weight and food intake in mice, manifested as a sustained decline in both body weight and cumulative food consumption. This has largely emphasized that dronedarone hydrochloride exhibits enormous potential in weight reduction and appetite suppression. When the difference in weight between the experimental and control groups of mice reached 10g, the modeling endpoint was reached. As we expected, the liver weight was significantly reduced, along with a marked decrease in fat content **(Figure 1D-I)**. It was worth noting that even when both the liver and fat mass were reduced, dronedarone hydrochloride still failed to decrease the content of skeletal muscle **(Figure 1J)**. Together, these findings indicate that dronedarone hydrochloride not only exhibits lipid-specific clearance in adipose tissue and the liver, but also has no adverse effects on skeletal muscle.

**Figure 1.**
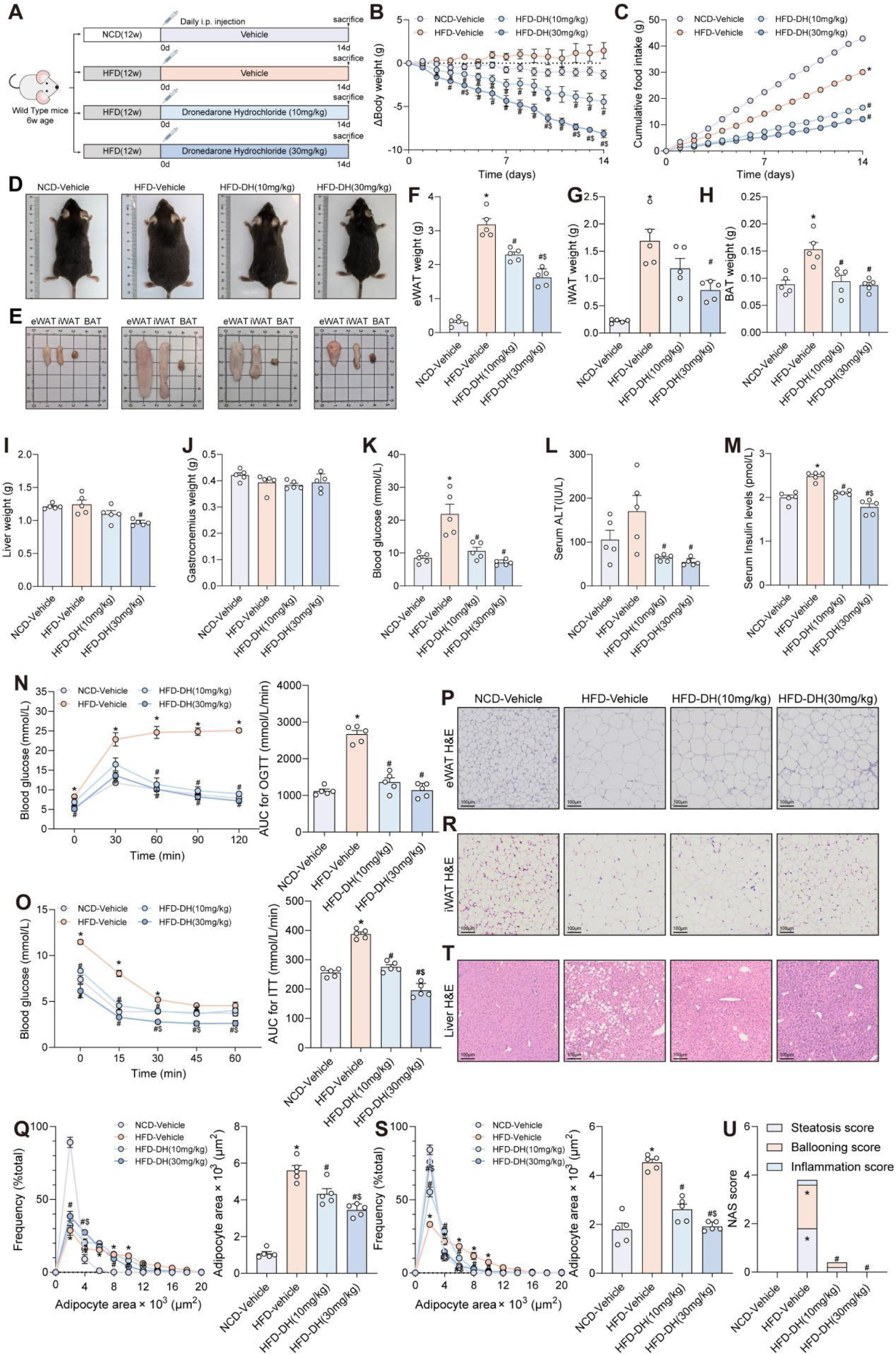
Dronedarone hydrochloride intraperitoneally to obese mice can treat obesity and improve obesity-related metabolic disorders. (A) The experimental diagram for the modeling of DH intraperitoneal administration in HFD-diet induced obesity (n=5 per group). (B-C) The daily changes of body weight and cumulative food intake in the process of Vehicle or DH administration. *p <0.05 vs. the NCD-Vehicle group; #p <0.05 vs. the HFD-Vehicle group; $p < 0.05 vs. the HFD-DH (10mg/kg) group (n=5 per group). (D-E) Representative images of general appearance for mice, eWAT, iWAT and BAT of Vehicle and DH-treated mice fed with NCD or HFD (n=5 per group). (F-j) Fat mass, liver and gastrocnemius weight among different groups. *p <0.05 vs. the NCD-Vehicle group; #p <0.05 vs. the HFD-Vehicle group (n=5 per group). (K-M) Serum blood glucose, ALT and insulin levels among different groups. *p <0.05 vs. the NCD-Vehicle group; #p <0.05 vs. the HFD-Vehicle group; $p < 0.05 vs. the HFD-DH (10mg/kg) group (n=5 per group). (N-O) Oral glucose tolerance test (OGTT), insulin tolerance test (ITT) and qualification for the area under curve after 14 days of Vehicle and DH administration. *p <0.05 vs. the NCD-Vehicle group; #p <0.05 vs. the HFD-Vehicle group (n=5 per group). (P-S) Hematoxylin and eosin (H&E) staining and qualification for the adipocyte frequency and area of eWAT and iWAT among different groups. *p <0.05 vs. the NCD-Vehicle group; #p <0.05 vs. the HFD-Vehicle group (n=5 per group). (T-U) Histological analyses of liver sections for Hematoxylin and eosin (H&E) staining and pathological scores (NAS score) assessing liver steatosis, ballooning and inflammation among different groups. *p <0.05 vs. the NCD-Vehicle group; #p <0.05 vs. the HFD-Vehicle group (n=5 per group).

As is well known, the accumulation of visceral fat triggers chronic low-grade inflammation and systemic insulin resistance, leading to a series of metabolic disorders involving glucose and lipid metabolism disorder and cardiovascular health. In the administration group, fasting blood glucose and serum ALT levels were all improved **(Figure 1K-L)**. Surprisingly, dronedarone hydrochloride also reduced insulin levels in obese mice **(Figure 1M)**. These findings suggest that dronedarone hydrochloride can improve pancreatic resistance and enhance pancreatic sensitivity. This hypothesis finding was also confirmed by OGTT and ITT tests in mice **(Figure 1N-O)**. Consistent with these observations, H&E staining results of eWAT and iWAT tissue sections demonstrated that dronedarone hydrochloride significantly inhibits lipid droplet formation and maturation **(Figure 1P-S)**. Liver H&E staining further revealed that dronedarone hydrochloride reduced lipid accumulation and inflammatory infiltration in the livers of mice **(Figure 1T-U)**. These findings uncover a novel dronedarone hydrochloride plays a crucial role in treating obesity, improving obesity-related metabolic disorders such as glucose and lipid dysregulation, and reducing inflammation.

### Oral administration of dronedarone hydrochloride can treat obesity and metabolic disorders in mice induced by a high-fat diet

After evaluating the anti-obesity effects of dronedarone hydrochloride observed with intraperitoneal injection, we applied oral gavage of dronedarone hydrochloride given that it is non-invasive and faithfully recapitulates human gastrointestinal absorption. C57BL/6J mice were randomly assigned into four groups and fed either with NCD or HFD for 12 weeks to establish a diet-induced obesity model and were later treated with vehicle, low-dose dronedarone hydrochloride (80mg/kg), or high-dose dronedarone hydrochloride (120mg/kg) by oral gavage daily for 13 days **(Figure 2A)**. Consistent with our earlier findings in intraperitoneal injection delivery, mice with dronedarone hydrochloride treatment exhibited significantly reduced body weight and cumulative food intake compared to their counterparts treated with vehicle in a dose dependent manner **(Figure 2B-C)**. Furthermore, gross morphology and assessment of organs’ weight further confirmed a leaner phenotype and decreased liver and adipose size in dronedarone hydrochloride-treated mice without affecting skeletal muscle mass **(Figure 2D-J)**.

**Figure 2.**
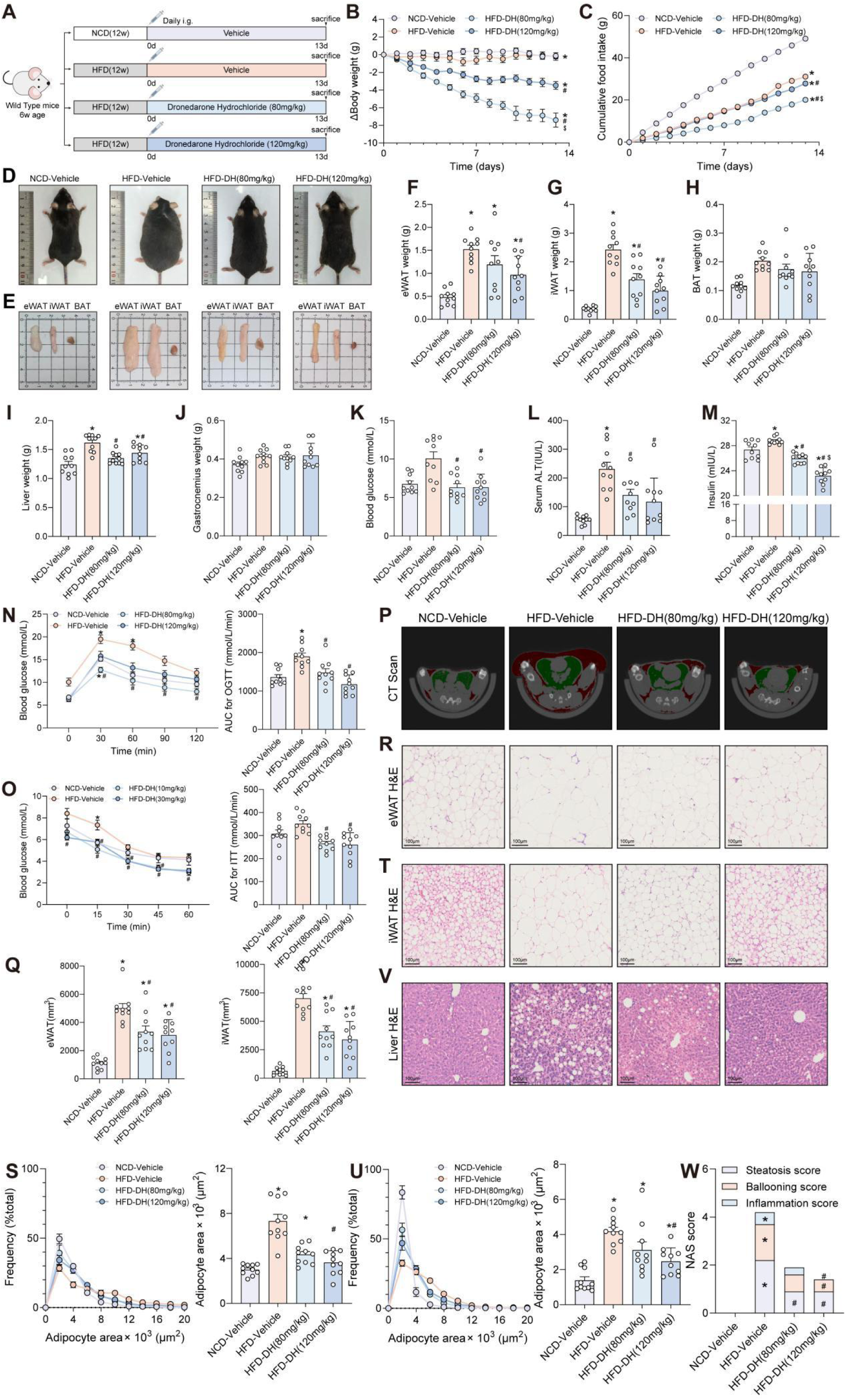
Gastric administration of decinatalone hydrochloride to obese mice can treat obesity and improve obesity-related metabolic disorders. (A) The experimental diagram for the modeling of DH oral administration in HFD-diet induced obesity (n=10 per group). (B-C) The daily changes of body weight and cumulative food intake in the process of Vehicle or DH administration. *p <0.05 vs. the NCD-Vehicle group; #p <0.05 vs. the HFD-Vehicle group; $p < 0.05 vs. the HFD-DH (80mg/kg) group (n=10 per group). (D-E) Representative images of general appearance for mice, eWAT, iWAT and BAT of Vehicle and DH-treated mice fed with NCD or HFD (n=10 per group). (F-J) Fat mass, liver and gastrocnemius weight among different groups. *p <0.05 vs. the NCD-Vehicle group; #p <0.05 vs. the HFD-Vehicle group (n=10 per group). (K-M) Serum blood glucose, ALT and insulin levels among different groups. *p <0.05 vs. the NCD-Vehicle group; #p <0.05 vs. the HFD-Vehicle group; $p < 0.05 vs. the HFD-DH (80mg/kg) group (n=10 per group). (N-O) Oral glucose tolerance test (OGTT), insulin tolerance test (ITT) and qualification for the area under curve after 13 days of Vehicle and DH administration. *p <0.05 vs. the NCD-Vehicle group; #p <0.05 vs. the HFD-Vehicle group (n=10 per group). (P-Q) Representative images of micro-CT analysis of mouse body composition and qualification of the eWAT and iWAT volume among different groups. *p <0.05 vs. the NCD-Vehicle group; #p <0.05 vs. the HFD-Vehicle group (n=10 per group). (R-U) Hematoxylin and eosin (H&E) staining and qualification for the adipocyte frequency and area of eWAT and iWAT among different groups. *p <0.05 vs. the NCD-Vehicle group; #p <0.05 vs. the HFD-Vehicle group (n=10 per group). (V-W) Histological analyses of liver sections for Hematoxylin and eosin (H&E) staining and pathological scores (NAS score) assessing liver steatosis, ballooning and inflammation among different groups. *p <0.05 vs. the NCD-Vehicle group; #p <0.05 vs. the HFD-Vehicle group (n=10 per group).

As anticipated, mice with oral gavage of dronedarone hydrochloride showed significantly lowered blood glucose and serum ALT levels compared to mice with vehicle **(Figure 2K–L)**, indicating simultaneous improvement in metabolic disorders associated with obesity involving hyperglycemia and hepatocellular injury. Serum insulin measurements revealed a marked attenuation of hyperinsulinemia and a restored insulin sensitivity in dronedarone hydrochloride-treated mice, according to the reduced area under the curve for both OGTT and ITT **(Figure 2M-O)**. In addition, to quantify depot-specific fat distribution, micro-CT based body composition analysis was performed. Both subcutaneous and visceral fat volumes were substantially decreased after dronedarone hydrochloride treatment **(Figure 2P-Q)**, providing a radiographic confirmation of adipose tissue reduction. Notably, histological analysis of adipose tissue demonstrated a significant reduction in adipocyte diameter and a corresponding increase in cell number per unit area, with quantitative analysis confirming reduced adipocyte size in both visceral and subcutaneous depots **(Figure 2R-U)**. Liver H&E staining further revealed dronedarone hydrochloride attenuated hepatic lipid accumulation and inflammatory infiltration in mice **(Figure 2V-W)**. Collectively, these findings demonstrate that oral dronedarone hydrochloride reverses obesity-induced insulin resistance and multi-organ metabolic dysfunction by inhibiting adipocyte hypertrophy and reducing hepatic lipid deposition, while preserving skeletal muscle mass, which is well suited to the treatment of obesity with metabolic syndrome.

### Dronedarone hydrochloride is comparable to semaglutide, tirzepatide and empagliflozin in terms of treatment efficacy

To position dronedarone hydrochloride within the current therapeutic landscape, we performed a head-to-head comparison with three FDA-approved first-line agents for metabolic disorders representing distinct mechanistic classes: Semaglutide, a GLP-1 receptor agonist; Tirzepatide, a GLP-1/GIP dual receptor agonist; and Empagliflozin, an SGLT-2 inhibitor. Mice fed a HFD were treated with vehicle, dronedarone hydrochloride (120 mg/kg, daily i.g. for 13 days), Semaglutide (3 nmol/kg, daily s.c. for 13 days) [16, 17], Tirzepatide (3 nmol/kg, daily s.c. for 13 days) [18], or Empagliflozin (20 mg/kg, daily i.g. for 13 days) [19] **(Figure 3A)**. Mice administered with the four drugs exhibited reduced body weight and cumulative food intake relative to vehicle group. Notably, dronedarone hydrochloride and Tirzepatide produced a comparable weight loss, both exceeding that achieved with Semaglutide or Empagliflozin **(Figure 3B-C)**. Similar reductions were observed in liver weight, iWAT, and eWAT tissue mass **(Figure 3D-I)**. Collectively, these findings demonstrate that dronedarone hydrochloride achieves weight reduction comparable to the first-line drugs.

**Figure 3.**
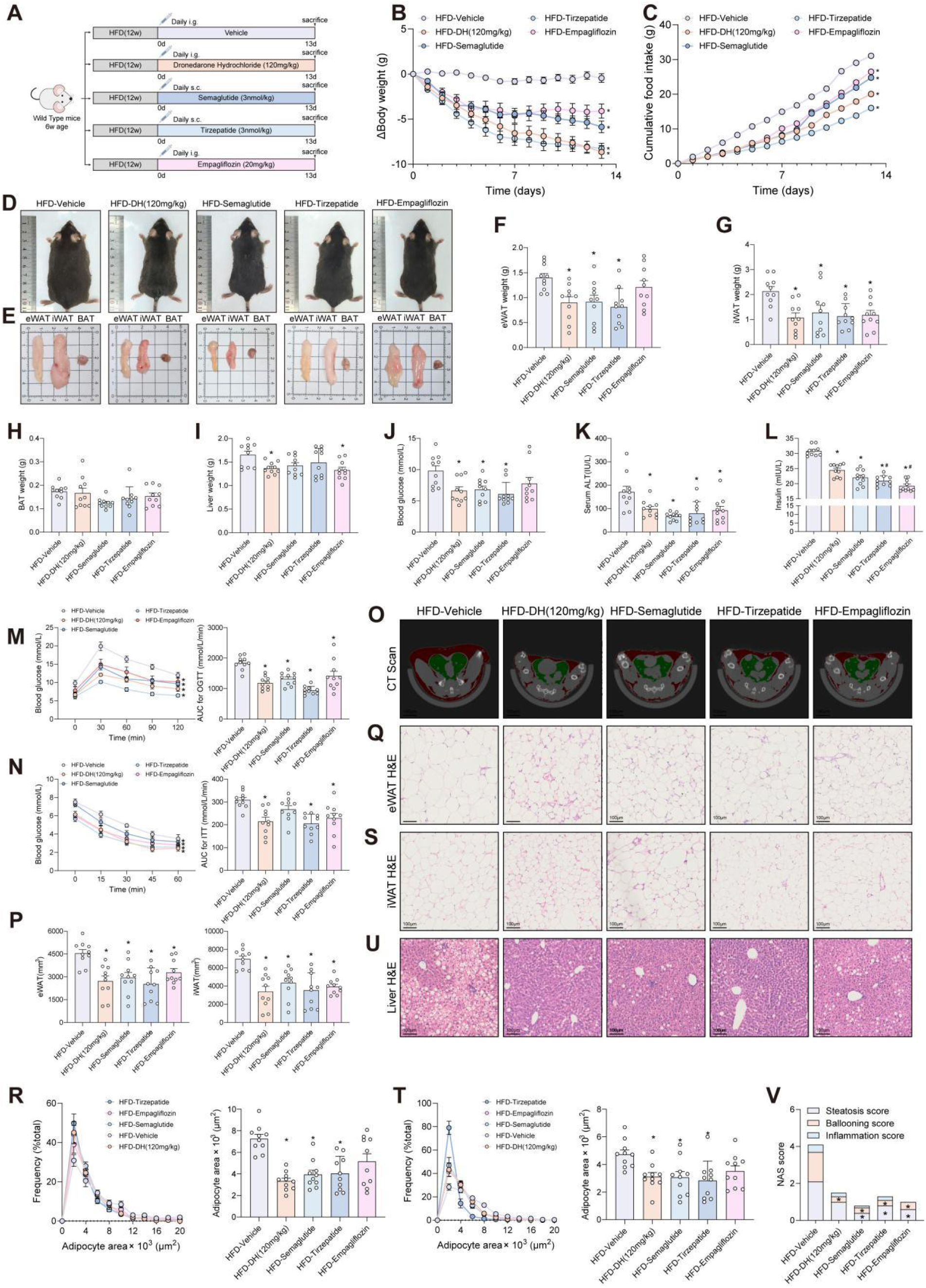
Dronedarone hydrochloride intervention is comparable to currently approved first-line agents. (A) The experimental diagram for the modeling of the effect comparison between DH and other first line drugs (Semaglutide, Tirzepatide and Empagliflozin) for obesity in HFD-diet induced obesity (n=10 per group). (B-C) The daily changes of body weight and cumulative food intake. *p <0.05 vs. the HFD-Vehicle group (n=10 per group). (D-E) Representative images of general appearance for mice, eWAT, iWAT and BAT of mice among different groups (n=10 per group). (F-I) Fat mass and liver weight among different groups. *p <0.05 vs. the HFD-Vehicle group (n=10 per group). (J-L) Serum blood glucose, ALT and insulin levels among different groups. *p <0.05 vs. the HFD-Vehicle group; #p <0.05 vs. the HFD-DH (120mg/kg) group (n=10 per group). (M-N) Oral glucose tolerance test (OGTT), insulin tolerance test (ITT) and qualification for the area under curve after 13 days of Vehicle or drug administration. *p <0.05 vs. the HFD-Vehicle group (n=10 per group). (O-P) Representative images of micro-CT analysis of mouse body composition and qualification of the eWAT and iWAT volume among different groups. *p <0.05 vs. the HFD-Vehicle group (n=10 per group). (Q-T) Hematoxylin and eosin (H&E) staining and qualification for the adipocyte frequency and area of eWAT and iWAT among different groups. *p <0.05 vs. the HFD-Vehicle group (n=10 per group). (U-V) Histological analyses of liver sections for Hematoxylin and eosin (H&E) staining and pathological scores (NAS score) assessing liver steatosis, ballooning and inflammation among different groups. *p <0.05 vs. the HFD-Vehicle group (n=10 per group).

We next measured whether dronedarone hydrochloride confers equivalent or superior benefits on the metabolic complications of obesity. All four agents significantly lowered blood glucose and serum ALT levels, and improved glucose tolerance and insulin sensitivity **(Figure 3J-L)**. OGTT and ITT analyses revealed that the glucose-lowering and insulin-sensitizing effects of dronedarone hydrochloride were non-inferior to the other three drugs **(Figure 3M-N)**. Micro-CT based body composition quantification confirmed that dronedarone hydrochloride reduced total body fat and visceral fat volume to a degree indistinguishable from the three positive comparators **(Figure 3O-P)**. Histologically, H&E staining of adipose tissue showed significantly reduced adipocyte size and lipid droplet area in all treatment arms, indicating suppression of pathological white adipose tissue expansion and the associated lipotoxicity, which was more apparent in epididymal adipose tissue **(Figure 3Q-T)**. Liver H&E staining revealed reduced hepatic lipid deposition and diminished inflammatory cell infiltration, confirming a reversal of hepatic steatosis and lobular inflammation **(Figure 3U-V)**. Collectively, these findings demonstrate that short-term dronedarone hydrochloride intervention achieves weight reduction, glycemic control, and hepatoprotection comparable to currently approved first-line agents, while attenuating visceral lipotoxicity and chronic systemic inflammation. These results support further development of dronedarone hydrochloride as a candidate for obesity-associated metabolic syndrome.

### Dronedarone hydrochloride does not affect the expression of appetite-regulating neuropeptides in the hypothalamus, either rely on the leptin pathway

The regulation of appetite and weight in humans is governed by a balance between anorexigenic and orexigenic signaling pathways, whereas obesity results from an imbalance between energy intake and expenditure [20]. The agouti-related peptide(AgRP)/neuropeptide Y(NPY) and pro-opiomelanocortin(POMC)/cocaine- and amphetamine-regulated transcript(CART) system are appetite-regulating peptides which both are located in the hypothalamic ARC, yet with their appetite-regulating peptide expression patterns being mutually exclusive and working in opposing directions [21]. Therefore, we measured the expression levels of four appetite peptides in mice to determine whether dronedarone hydrochloride’s effect on mouse appetite is mediated by regulation of the hypothalamic appetite peptide system. Strikingly, the experimental results revealed that dronedarone hydrochloride does not affect the expression of appetite-regulating neuropeptides in the hypothalamus **(Figure 4A-D)**. The hormone leptin also plays a role in food intake, body mass, proinflammatory immune responses, angiogenesis and lipolysis. We subsequently used dronedarone hydrochloride to treat spontaneous obesity in ob/ob mice induced by leptin deficiency, in order to investigate whether the action of dronedarone hydrochloride was influenced by leptin **(Figure 4E)**. We observed that no matter the improvements in body weight and food intake or the reductions in liver weight and fat mass, were consistent with previous findings **(Figure 4F-M)**. Dronedarone hydrochloride could still reduce body weight, decrease food intake, and improve obesity even in the absence of leptin. Furthermore, dronedarone hydrochloride did not affect muscle mass while promoting weight loss **(Figure 4N)**. Compared with the control group, ob/ob mice treated with dronedarone hydrochloride also showed great improvements in blood glucose levels and glucose regulation. Measurement of fasting blood glucose levels in mice confirmed that dronedarone hydrochloride could significantly alleviate hyperglycemia **(Figure 4O)**. Following drug intervention, the AUC of blood glucose levels in both OGTT and ITT tests decreased simultaneously, indicating that dronedarone hydrochloride can effectively improve glucose tolerance, enhance systemic insulin sensitivity, significantly alleviate insulin resistance secondary to obesity, and comprehensively correct glucose metabolism abnormalities **(Figure 4P-Q)**. These findings uncover a novel that the therapeutic effect of dronedarone hydrochloride in obesity-related metabolic syndrome does not rely on the leptin pathway.

**Figure 4.**
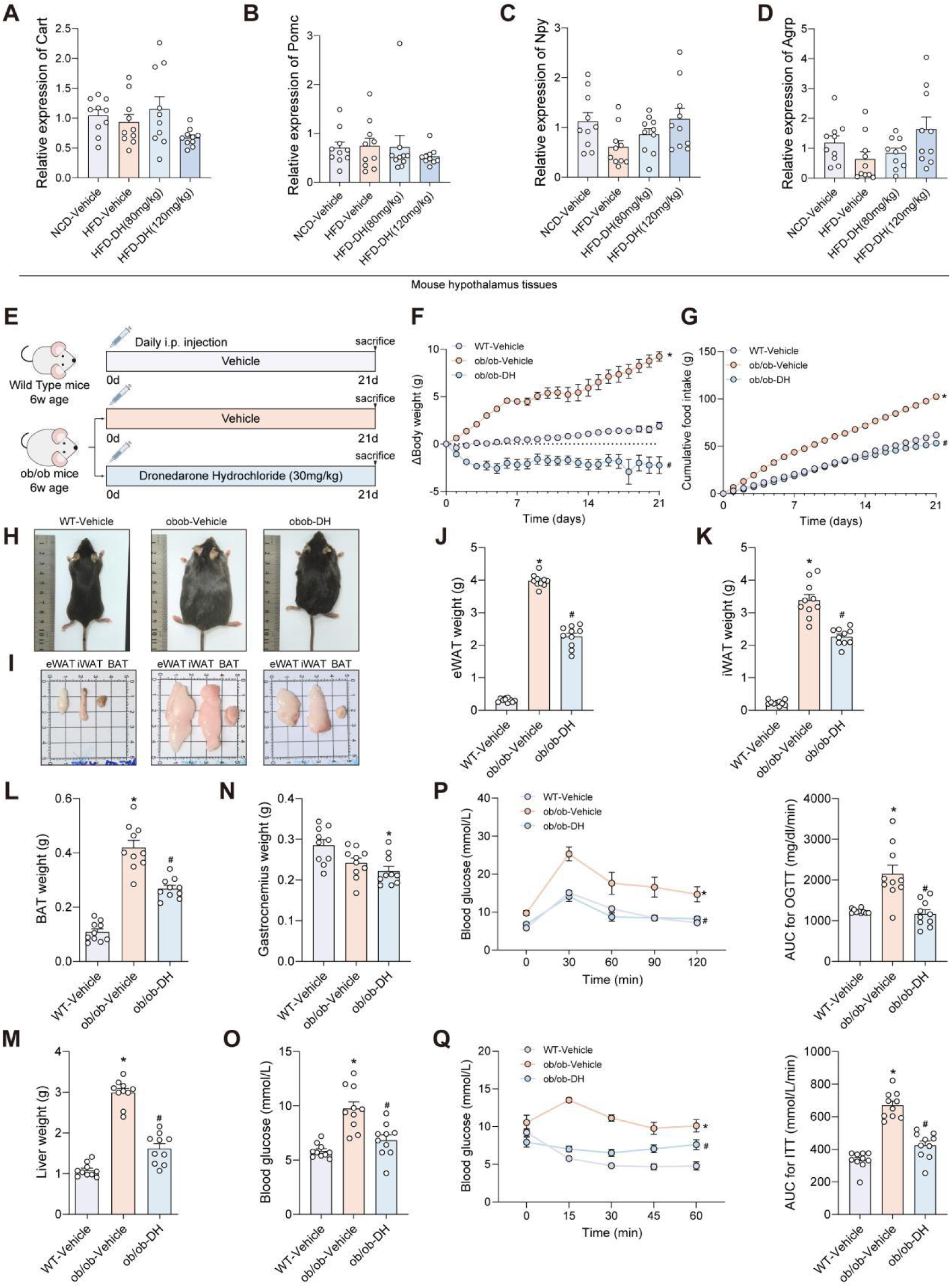
The therapeutic effect of dronedarone hydrochloride does not rely on appetite-regulating neuropeptides in the hypothalamus and the leptin pathway. (A-D) Expression mRNA of appetite-regulating peptide expression (Agrp/Npy and Pomc/Cart) in the hypothalamic ARC of the NCD-Vehicle group, HFD-Vehicle group and the HFD-DH (80mg/kg, 120mg/kg) group (n=10 per group). (E) The experimental diagram for the modeling of DH intraperitoneal injection in ob/ob mice (n=10 per group). (F-G) The daily changes of body weight and cumulative food intake in the process of Vehicle or DH administration. *p <0.05 vs. the WT-Vehicle group; #p <0.05 vs. the ob/ob-Vehicle group (n=10 per group). (H-I) Representative images of general appearance for mice, eWAT, iWAT and BAT of Vehicle and DH-treated mice (n=10 per group). (J-N) Fat mass, liver and gastrocnemius weight among different groups. *p <0.05 vs. the WT-Vehicle group; #p <0.05 vs. the ob/ob-Vehicle group (n=10 per group). (O) Serum blood glucose levels among different groups. *p <0.05 vs. the NCD-Vehicle group (n=10 per group). (P-Q) Oral glucose tolerance test (OGTT), insulin tolerance test (ITT) and qualification for the area under curve after 21 days of Vehicle and DH administration. *p <0.05 vs. the NCD-Vehicle group; #p <0.05 vs. the HFD-Vehicle group (n=10 per group).

## DISCUSSION

The therapeutic effect of dronedarone hydrochloride in obesity-related metabolic syndrome has been described in this study, corroborating the data observed here. Indeed, C57BL/6 mice fed HFD for 12 weeks and later treated with vehicle, low-dose or high-dose dronedarone hydrochloride experienced significant body weight loss. The two-week drug intervention significantly reduced food intake, the weight of the liver, subcutaneous and visceral fat pads. Micro-CT results confirmed that the drug significantly reduces both total systemic fat and visceral fat accumulation. We also demonstrated that dronedarone hydrochloride reduces fasting blood glucose and insulin levels, decreases the area under the OGTT and ITT curves, and simultaneously lowers serum total cholesterol and ALT levels. Moreover, there is evidence that dronedarone hydrochloride reduces the accumulation of lipid droplets in hepatocytes, alleviates inflammatory infiltration within the liver, and reduces the volume of fat cells, thereby effectively alleviating lipotoxicity and systemic chronic low-grade inflammation. Building on this evidence, we propose that dronedarone hydrochloride can improve obesity, dyslipidaemia and hepatic steatosis, and exert a distinct therapeutic effect on obesity complicated by insulin resistance, which offers novel mechanistic insights into the clinical treatment of obesity-related metabolic disorders.

This study conducted comparisons using three first-line metabolic drugs, semaglutide, tirzopentide and empagliflozin as the positive controls. These results showed that dronedarone hydrochloride combined efficacy in terms of weight loss, fat reduction, and improvement in glucose tolerance and insulin sensitivity was non-inferior to that of these three marketed drugs. Semaglutide, a GLP-1 receptor agonist and Tirzepatide, a GLP-1/GIP dual receptor agonist, exert their effects by suppressing appetite and delaying gastric emptying through central nervous system inhibition. Empagliflozin, an SGLT-2 inhibitor, lowers blood glucose by promoting urinary glucose excretion via renal SGLT-2-mediated transport [22]. However, all of them exhibit inherent defects. For example, weight loss induced by GLP-1 receptor agonists is accompanied by skeletal muscle wasting, which can easily lead to a reduction in basal metabolic rate and a rebound effect upon discontinuation [23]. Moreover, the lipid-lowering effect of Empagliflozin is relatively limited [24]. By comparison, dronedarone hydrochloride achieves equivalent effects in terms of weight loss, blood glucose control, and liver protection, whilst fully preserving skeletal muscle mass. Our findings carry important translational potential for obesity treatment among middle-aged and elderly individuals at high risk for obesity, fatty liver disease, and sarcopenia.

Nevertheless, this study has room for further exploration. We systematically verified the pharmacodynamic effects of dronedarone hydrochloride, including weight loss, hypoglycemia, liver protection and muscle preservation. Still, its precise molecular mechanisms remain unclear. This work did not identify the key targets and signaling pathways mediating its regulation of lipid metabolism and insulin sensitivity. Future studies will conduct molecular and cellular experiments to clarify its mechanisms.

We uncovered that short-term dronedarone hydrochloride therapy achieves weight loss and lipid reduction effects comparable to those of semaglutide, tiramalide, and empagliflozin, while improving glucose-lipid metabolism and mitigating liver injury. Additionally, it alleviates tissue inflammation and reverses obesity-related insulin resistance by reducing visceral lipid deposition. Compared to existing first-line clinical drugs, the most pivotal innovative feature of dronedarone hydrochloride is its ability to reduce fat accumulation while preserving skeletal muscle mass, addressing the clinical limitations of GLP-1 receptor agonists in causing muscle loss and SGLT-2 inhibitor in exhibiting weak lipid-lowering efficacy, thereby providing more sustained maintenance of metabolic homeostasis. Although the specific molecular regulatory mechanisms still require further investigation, the existing body of evidence, encompassing imaging, biochemistry, functional studies and histopathology, has fully demonstrated the benefits of this comprehensive metabolic intervention. It provides a novel interventional candidate strategy for obesity-related metabolic syndrome, with promising prospects for clinical translation and new drug development.

## ACKNOWLEDGMENTS

We thank our team members (Cholestatic Liver Diseases Center and Metabolic Dysfunction-Associated Steatotic Liver Disease (MASLD) Medical Research Center, The First Affiliated Hospital (Southwest Hospital) of Third Military Medical University (Army Medical University) for their generous help.

